## extended materials and reagents for "Antibody-directed extracellular proximity biotinylation reveals Contactin-1 regulates axo-axonic innervation of axon initial segments"

Reagents, antibodies, and plasmids used in this study are summarized below:

| REAGENT or RESOURCE | SOURCE | IDENTIFIER |
|---|---|---|
| **Antibodies** | | |
| Mouse anti-AnkG (IgG2a) | NeuroMab | Cat# 75-146; RRID: AB_10673030 |
| Mouse anti-Gephyrin (IgG1) | Synaptic Systems | Cat# 147011; RRID: AB_887717 |
| Guinea pig anti-VGAT | Synaptic Systems | Cat# 131004; RRID: AB_887873 |
| Rabbit monoclonal anti-HA | Cell Signaling Technology | Cat# 3724; RRID: AB_1549585 |
| Mouse anti-Myc | MBL International Corporation | Cat# M192 PRID: AB_11160947 |
| Mouse anti-PSD95 | Antibodies Incorporated | Cat# 75-028 PRID: AB_2292909 |
| Mouse anti-Tenascin-R | R&D Systems | Cat# MAB1624 RRID: AB_2207001 |
| Mouse anti-Tuj1 | BioLegend | Cat# 801202 RRID: AB_10063408 |
| Mouse anti-V5 | Invitrogen | Cat# R960CUS RRID: AB_159298 |
| Rabbit anti-βIV-Spectrin | Rasband lab | RRID: AB_2315634 |
| Rabbit anti-Kv1.2 | Gift of Dr. James Trimmer | PRID: 2756300 |
| Rabbit anti-NrCAM | Abcam | Cat# ab24344 RRID: AB_448024 |
| Chicken anti-MAP2 | Encor | Cat# CPCA-MAP2 |
| Chicken anti-Neurofascin | R&D Systems | Cat# AF3235 RRID: AB_10890736 |
| Goat anti-Cntn1 | R&D Systems | Cat# AF904 RRID: AB_2292070 |
| Rat anti-HA | Millipore Sigma | Cat# 11867423001 RRID: AB_390918 |

| | | |
|---|---|---|
| HRP-conjugated goat anti-chicken IgY | Aves Labs | Cat# H-1004<br>RRID: AB_2313517 |
| HRP-conjugated goat anti-rabbit IgG | Jackson ImmunoResearch Labs | Cat# 111-035-003<br>PRID: AB_2313567 |
| Alexa Fluor 594 conjugated streptavidin | Thermo Fisher Scientific | Cat# S11227 |
| Aminomethylcoumarin (AMCA) conjugated goat anti-chicken IgY | Jackson ImmunoResearch Labs | Cat# 103-155-155<br>RRID: AB_2337385 |
| Aminomethylcoumarin (AMCA) conjugated goat anti-rat IgG | Thermo Fisher Scientific | Cat# A21093<br>PRID: AB_2535748 |
| Alexa Fluor 488 conjugated goat anti-chicken IgY | Jackson ImmunoResearch Labs | Cat# 103-545-155<br>RRID: AB_2337390 |
| Alexa Fluor 488 conjugated goat anti-mouse IgG | Thermo Fisher Scientific | Cat# A11029<br>RRID: AB_2534088 |
| Alexa Fluor 488 conjugated goat anti-rabit IgG | Thermo Fisher Scientific | Cat# A11034<br>RRID: AB_2758380 |
| Alexa Fluor Plus 594 conjugated goat anti-mouse IgG | Thermo Fisher Scientific | Cat# A32742<br>RRID: AB_2762825 |
| Alexa Fluor 594 conjugated donkey anti-goat IgG | Thermo Fisher Scientific | Cat# A11058<br>RRID: AB_2758385 |
| Alexa Fluor 488 conjugated goat anti-rat | Thermo Fisher Scientific | Cat# A-11006; RRID: AB_2534074 |
| Alexa Fluor Plus 555 conjugated goat anti-rabbit | Thermo Fisher Scientific | Cat# A32732; RRID: AB_2633281 |
| Alexa Fluor 647 conjugated goat anti-rabbit | Thermo Fisher Scientific | Cat# A-21244; RRID: AB_2535812 |
| Alexa Fluor 555 conjugated goat anti-guinea pig | Thermo Fisher Scientific | Cat# A-21435; RRID: AB_2535856 |
| Alexa Fluor 647 conjugated goat anti-guinea pig | Thermo Fisher Scientific | Cat# A-21450; RRID: AB_2735091 |
| Alexa Fluor 647 conjugated goat anti-mouse IgG1 | Thermo Fisher Scientific | Cat# A-21240; RRID: AB_2535809 |
| Alexa Fluor 488 conjugated goat anti-mouse IgG2a | Thermo Fisher Scientific | Cat# A-21131; RRID: AB_2535771 |
| **Bacterial and Virus Strains** | | |
| Stellar competent cells | Takara | Cat# 636763 |

| | | |
|---|---|---|
| NEB® Stable Competent E. coli (High Efficiency) | New England Biolabs | Cat# C3040H |
| NEB® 5-alpha Competent E. coli (High Efficiency) | New England Biolabs | Cat# C2987H |
| AAV PHP.S encapsulated vectors | This paper | N/A |
| AAV PHP.eB encapsulated vectors | This paper | N/A |
| **Chemicals, Peptides, and Recombinant Proteins** | | |
| Biotin tyramide | Perkin Elmer | Cat# NEL749A001KT |
| Neurobasal Medium | Gibco | Cat# 21103-049 |
| B-27 supplement | Gibco | Cat# 17504-044 |
| GlutaMax-I | Gibco | Cat# 35050-061 |
| Penicillin-Streptomycin | Gibco | Cat# 15140122 |
| DMEM, high glucose, no glutamine | Gibco | Cat# 11960-044 |
| Fetal bovine serum | Cytiva | Cat# SH30109.03 |
| 0.5% Trypsin-EDTA | Gibco | Cat# 15400054 |
| 2.5% trypsin | Gibco | Cat# 15090046 |
| Hank's buffered salt solution (HBSS) | Gibco | Cat# 14175-095 |
| Poly-D-lysine | Gibco | Cat# A3890401 |
| Poly-L-lysine | Sigma | Cat# P4832-50ML |
| Laminin | Gibco | Cat# 23017015 |
| Phosphate buffer saline (PBS) | Corning | Cat# 21-031-CV |
| Opti-MEM | Thermo Fisher Scientific | Cat# 31985070 |
| Hoechst 33258 | Thermo Fisher Scientific | Cat# H3569<br>RRID: AB_2651133 |
| Polyethylenimine (PEI) | Polysciences | Cat# 24765-1 |
| Normal goat serum | Gibco | Cat# 16210-072 |
| Gelatin from bovine skin | Sigma | G9382-100G |
| FastGreen | Sigma | Cat# F7258 |
| Streptavidin Mag Sepharose | GE Healthcare | Cat# 28-9857-38 |
| Vectashield plus anti-fade mounting media | Vector labs | Cat# H-1900 |
| Sequencing Grade Trypsin | Promega | V5111 |
| Tamoxifen | Sigma-Aldrich | Cat# T5648; CAS:<br>10540-29-1 |
| Fluoromount-G | Southern Biotech | 0100-01 |
| **Critical Commercial Assays** | | |
| Q5 High-Fidelity DNA Polymerase | New England Biolabs | Cat# M0491S |

| | | |
|---|---|---|
| CloneAmp HiFi PCR Premix | Takara | Cat# 639298 |
| NucleoSpin Plasmid Transfection-grade | Takara | Cat# 740490.250 |
| In-Fusion Snap Assembly Master Mix | Takara | Cat# 638947 |
| Gel purification | Cytiva | 28903470 |
| Amicon Ultra-15 Centrifugal Filter Unit 100 kDa | Millipore | Cat# UFC910008 |
| C18 ZipTips | Millipore | Cat# ZTC18S096 |
| BamHI restriction enzyme | New England Biolabs | Cat# R3136T |
| EcoRI restriction enzyme | New England Biolabs | Cat# R3101T |
| NotI restriction enzyme | New England Biolabs | Cat# R3189L |
| XhoI restriction enzyme | New England Biolabs | Cat# R0146M |
| **Experimental models: Cell Lines** | | |
| HEK293T cells | Provided from Baylor College of Medicine Neuroconnectivity Core | N/A |
| **Experimental models: Organisms/ Strains** | | |
| Rat: Sprague-Dawley rat embryos | Charles River Laboratories | SAS-SD |
| Mouse: C57BL/6 | Baylor College of Medicine Center for Comparative Medicine | N/A |
| Mouse: CFW (Swiss Webster) | Charles River | Cat# CRL:24; RRID: IMSR_CRL:24 |
| Mouse: Nkx2-1$^{tm1.1(Cre/ERT2)Zjh}$/J | Gift from Z.J. Huang; (Taniguchi et al., 2013) | JAX: 014552; RRID: IMSR_JAX:014552 |
| Mouse: B6;129S6-Gt(Rosa)26Sor$^{tm9(CAG-tdTomato)Hze}$/J | Gift from Z.J. Huang; (Taniguchi et al., 2013) | JAX: 007905; RRID: IMSR_JAX:007905 |
| Mouse: Igs2tm1.1(CAG-cas9*)Mmw/J (also known as H11Cas9 mice) | The Jackson Laboratory | Cat# JAX: 027650 RRID: IMSR_JAX:027650 |
| Mouse: Cntn1 knockout | Boyle et al., 2001 | Cat# JAX: q034216, RRID: IMSR_JAX:034216 |
| Mouse: ICR | Baylor College of Medicine Center for Comparative Medicine | N/A |

| Oligonucleotides | |
|---|---|
| **Target for knock-in** | **Sequence (5'->3')** |
| rat Adgrl1 #1 | TTGAGGGGCCAGGGCCCGAT |
| rat Adgrl1 #2 | GTTGGTCACTAGTCTCTGAG |
| rat Cacna2d1 #1 | ATACTCCTTTGGCTGGTATC |
| rat Cacna2d1 #2 | CTGGCAGCAGACACTATCTA |
| rat Celsr2 #1 | CGCCGGCCCAGGGTTAATGC |
| rat Celsr2 #2 | ATCGCCATGAGCATCAAGGC |
| rat Cntn1 #1 | GGTGTGTCCACGCTGTCATC |
| rat Cntn1 #2 | GAGGAGACCCGATGACAGCG |
| rat Dcc #1 | GTTTGGCAATGCATGTCAAA |
| rat Dcc #2 | AAGCAACTCAATGCCATCAC |
| rat Fat3 #1 | CTACGGGTGTCCTCTACACC |
| rat Fat3 #2 | GGAAACGCAGCACCAGACCC |
| rat Kctd12 #1 | ACACCGAGTACGTCTTCTGC |
| rat Kctd12 #2 | CTCCCTGCAGAAGACGTACT |
| rat L1Cam #1 | GCAGTAGCCCTAGAATAGCA |
| rat L1Cam #2 | GACTGGACCTTGCTATTCTA |
| rat Lrp1 #1 | ATAGGGGATCCCTTAGCATA |
| rat Lrp1 #2 | TGGGGCAGGGCCCTATGCTA |
| rat Lrrc4b #1 | GCTCTTCAAGAGCGGCTCCA |
| rat Lrrc4b #2 | GAGCCGCTCTTGAAGAGCAG |
| rat Ncam1_v1,2,5 #1 | GGGACCGTCTTGACTTCAGT |
| rat Ncam1_v1,2,5 #2 | CTCATTCTCTTTCGTTTGTG |
| rat Ncam1_v3,4 #1 | AGGGTGGCTGGGATGGCCGT |
| rat Ncam1_v3,4 #2 | CAATGAGACCAAGGTGTAGG |
| rat Neo1 #1 | AGGTCATCAAGCCGTTGTGA |
| rat Neo1 #2 | GGACCTAAATGCCATCACAA |
| rat Nfasc #1 | AGTCAATGCCATCTATTCCC |
| rat Nfasc #2 | CCATCTATTCCCTGGCCTGA |
| rat Nlgn3 #1 | ATAGAGCTATACACGGGTAG |
| rat Nlgn3 #2 | TCTGGGATAGAGCTATACAC |
| rat NrCAM #1 | AAGGAGTTCATTGCGTTGAC |
| rat NrCAM #2 | GAGCTTAAAGACTTAAACAA |

| | |
|---|---|
| rat Nrxn1 #1 | GTCCAATGGGGCTGTGGTCA |
| rat Nrxn1 #2 | CTCCTTGACCACAGCCCCAT |
| rat Plxna2 #1 | GCAACGACTGGCCTACAAGG |
| rat Plxna2 #2 | CATGTCCATAGAGAGCTGAA |
| rat Plxna4 #1 | CACACTGGTCATCATGGTCC |
| rat Plxna4 #2 | TCAGCTGTCTAAGCTCATGA |
| rat Plxnb2 #1 | AACCTTATTCTCGAGCGCAG |
| rat Plxnb2 #2 | TGCCGCTGCGCTCGAGAATA |
| rat Ptprs #1 | CCAGGTATTCCAACGCCGCC |
| rat Ptprs #2 | GTTCTGCTTCCAGGCGGCGT |
| rat Tenm4 #1 | GACAGAGCGAGATGGGCCGA |
| rat Tenm4 #2 | TTCATGAGACAGAGCGAGAT |
| rat Cntn2 #1 | GGAGAACGCAGGTCCGGGAT |
| rat Cntn2 #2 | ATGGTGATATTGATGCTCGC |
| mouse Cntn1 #1 | GAGGAGACTCGAAGACAGCG |
| mouse Cntn1 #2 | CCTTGGCTTTCTTGTCTACT |
| mouse Cntn1 #3 | CGAGTAGACAAGAAAGCCAA |

| **Target for knock-out** | **Sequence (5'->3')** | **Sequence (5'->3')** | **Sequence (5'->3')** |
|---|---|---|---|
| Control | CTGTCTTCATCATGGCCGAC | GTTCGCATTATCCGAACCAT | TAAGCGTCGCAAGAAGACGG |
| Nfasc | ACGATCTCGGTGAGAGTAAA | CATGTCCTGCAGCATGACGT | ACTGCGGAAGTCGATCACCA |
| Cntn1 | GCATAGCAGAAAACGCGTAT | CTACGTGATTGACTTTAACA | TCGGATGAGCAGTTCCCCAT |
| NrCAM | ATGTGTTCCGTTACGAGTCC | GCCGGTGGTAATCCAATTGG | GGATTACAATATTCTCCCGT |

| **Recombinant DNA plasmids** | | |
|---|---|---|
| PX551 (AAV-SpCas9) | Gift of Dr. Feng Zhang | Addgene #60957 |
| PX552 (pAAV-U6sgRNA(SapI)_hSyn-GFP-KASH-bGH) | Gift of Dr. Feng Zhang | Addgene #60958 |
| pCAG-1BPNLS-Cas9-1BPNLS | Gift of Dr. Juan Belmonte | Addgene #87108 |
| pCAG_smFP HA | Gift of Dr. Loren Looger | Addgene #59759 |
| pCAG_smFP V5 | Gift of Dr. Loren Looger | Addgene #59758 |
| pMJ114 | Gift of Dr. Jonathan Weissman | Addgene #85995 |
| pMJ117 | Gift of Dr. Jonathan Weissman | Addgene #85997 |
| pMJ179 | Gift of Dr. Jonathan Weissman | Addgene #85996 |
| pUCmini-iCAP-PHP.S | Gift of Dr. Viviana Gradinaru | Addgene #103002 |
| pUCmini-iCAP-PHP.eB | Gift of Dr. Viviana Gradinaru | Addgene #103005 |

| | | |
|---|---|---|
| pHelper | Agilent Technologies | Cat #240071 |
| Knock-in sgRNA and donor plasmid | This paper | N/A |
| pAAV-3x-gRNA-smFP | This paper | N/A |
| rat contactin-myc | Gift of Dr. Elior Peles | N/A |
| pAAV-hSyn-EGFP | Gift of Dr. Bryan Roth | Addgene # 5046 |
| pcDNA3 | Invitrogen | N/A |
| hSYNp-Cntn1-Myc full-length | This paper | N/A |
| hSYNp-Cntn1-Myc truncates | This paper | N/A |
| **Software and Algorithms** | | |
| Fiji | NIH | https://fiji.sc/ |
| Zeiss Zen (Blue Edition) Imaging Software | Zeiss | http://www.zeiss.com/microscopy/en_us/products/microscope-software/zen.html#introduction; RRID: SCR_013672 |
| NIS Elements Imaging Software | Nikon | https://www.microscope.healthcare.nikon.com/products/software/nis-elements |
| CRISPOR | Haeussler et al., 2016 | http://crispor.tefor.net/crispor.py |
| Adobe Photoshop (version 23.5.1) | Adobe | https://www.adobe.com/products/photoshop.html |
| Adobe Illustrator (version 26.5) | Adobe | https://www.adobe.com/products/illustrator.html |
| GraphPad Prism 9 | GraphPad | https://www.graphpad.com/ |
| MATLAB | Mathworks | https://www.mathworks.com/products/matlab.html |
| Python | Python Software Foundation | https://www.python.org/ |

| | | |
|---|---|---|
| Script for calculation of extracellular tyrosines | Rasband lab | https://github.com/jrasband/extracellular-tyrosines |
| BioRender | BioRender | https://biorender.com/ |
| **Other** | | |
| Axioimager Z2 with Apotome | Zeiss | |
| Ni2 upright fluorescence microscope | Nikon | |
| Microtome HM450 | Thermo Scientific | |
| Cryostat CryoStar NX70 | Thermo Scientific | |
| QExactive Plus | Thermo Scientific | |
| NanoAcquity Ultra Performance UPLC system | Waters | |
| 15-cm EasySpray C18 column | Thermo Scientific | |
